# Keeping patient phenotypes and genotypes private while seeking disease diagnoses

**DOI:** 10.1101/746230

**Authors:** Karthik A. Jagadeesh, David J. Wu, Johannes A. Birgmeier, Dan Boneh, Gill Bejerano

## Abstract

In an age where commercial entities are allowed to collect and directly profit from large amounts of private information, an age where large data breaches of such organizations are discovered every month, science must strive to offer society viable ways to preserve privacy while benefitting from the power of data sharing. Patient phenotypes and genotypes are critical for building groups of phenotypically-similar patients, identify the gene that best explains their common phenotypes, and ultimately, diagnose a patient with a Mendelian disease. Direct computation over these quantities requires highly-sensitive patient data to be shared openly, compromising patient privacy and opening patients up for discrimination. Existing protocols focus on secure computation over genotype data and only address the final steps of the disease-diagnosis pipeline where phenotypically-similar patients have been identified. However, identifying such patients in a secure and private manner remains open. In this work, we develop secure protocols to maintain patient privacy while computing meaningful operations over *both* genotypic and phenotypic data for two real scenarios: COHORT DISCOVERY and GENE PRIORITIZATION. Our protocols newly enable a complete and secure *end-to-end disease diagnosis pipeline* that protects sensitive patient phenotypic and genotypic data.

## Introduction

We can now accurately diagnose over 5,000 monogenic diseases and attribute their causes to over 3,400 different disease genes(*1*). Each year, approximately 300 novel monogenic disease genes are discovered(*2*); revealing novel disease mechanisms and disease genes(*3*–*5*). Obtaining a precise disease diagnosis enables better treatment plans, disease management, and care for the patient(*6, 7*). The diagnosis also provides a sense of closure to the patient family, informs family counseling, and in the age of genome editing, provide first hope for a cure.

However, to identify the *single* gene that causes a patient’s disease, a patient must reveal their entire genome to the test provider together with their full set of ailments, signs and symptoms(*8*). This highly personal information can potentially be used to discriminate against not only the patient themselves, but also their next of kin. At the same time, over 99.9% of this sensitive patient information is completely *irrelevant* to a monogenic disease diagnosis (and in principle, need *not* be shared at all).

Previously, we have shown(*9*) how exact computations on genomic data can be performed without sharing genomic inputs. For instance, a small cohort of strangers suspected of having the same, yet-to-be-diagnosed disease, can discover whether they possess a mutation in the same gene without sharing any other genomic data with each other or with the test provider. In a similar vein, our techniques can identify the causal mutation present in affected family members but not in unaffected relatives (e.g., cousins), without any family member needing to reveal their genome to each other or to the test provider. In both scenarios, virtually nothing is revealed about the genomes of the participants, and moreover, the precise output of the analysis is successfully computed.

In an era of great genomic discoveries, but also great concerns for privacy, and great data breaches, our ability to provide private genomic services without revealing participant genomes must be extended. While the secure protocols over patient genomes described above are critical for obtaining a diagnosis, obtaining the necessary inputs to those algorithms typically requires computing over the (equally-sensitive) patient phenotype data. For instance, several of the tests require identifying cohorts of phenotypically-similar patients; it is not clear how to establish such cohorts without having patients collectively share and compare their phenotypes. In this work, we extend the scope of patient privacy beyond protecting genomic data to additionally protect patient *phenotypic data*. Our work newly enables privacy-preserving versions of the following exact tests:

First, a large group of undiagnosed strangers can come together and determine whether any two or more of them share a promising set of phenotypes without needing to reveal any of their disease phenotypes to each other or to the test provider. If one or more such small groups are found, each small cohort can then rely on our previously-developed privacy-preserving protocols to identify a common-mutated gene that explains their phenotypes without needing to share their genomic information(*9*). For those patients for whom a small cohort is not identified (often a majority), virtually *nothing* is revealed about their own condition.

A full 70% of patients sequenced for a monogenic disease cannot be immediately diagnosed (in part because the causative gene for the disease is currently unknown), and can greatly benefit from the above service(*10, 11*). For the remaining 30% of patients on average where a diagnosis is possible, they would benefit from consulting a commercial entity that specializes in monogenic disease diagnosis to obtain the actual diagnosis based on their set of candidate disease genes and observed phenotypes. In our second advancement, we design a protocol that enables a patient to consult with the commercial service provider and learn a handful of potentially-causal genes for their condition without needing to reveal any of their genotype or phenotype information to the commercial service provider. Moreover, the service provider only reveals a sliver of their competitive knowledge by only revealing to the patient a shortlist of potentially-causal genes from their much larger list of candidate disease genes.

## Results

### Quantifying the similarity of any two sets of phenotypes

Virtually no two patients suffering from the same monogenic disease will be described using identical phenotypic terms. They may exhibit somewhat different phenotypes or there could be differences in the granularity of terms used by their different clinicians, differences in the battery of tests they have undergone or the symptoms they choose to share with their clinicians. The Human Phenotype Ontology(*12*) (HPO) is extremely helpful as it organizes sets of phenotypes into a structured hierarchy.

HPO is a structured vocabulary (technically, a Directed Acyclic Graph, or DAG) that attempts to capture all genetically-derived human phenotypes in the form of a hierarchy. For example, the HPO term “Hypomimic face” is a child term of both “Decreased facial expression” and “Abnormality of facial musculature,” and the latter is in turn a child of “Abnormality of facial soft tissue” and so on.

Recently, we developed the Phrank (for PHenotype RANKing) algorithm for quantifying the similarity between any two sets of HPO phenotypes, and showed that Phrank improves on the performance of previous algorithms in tasks like obtaining a differential genetic disease diagnosis based on a set of patient phenotypes and a knowledgebase of gene-phenotype-disease associations(*13*). Here we develop a privacy-preserving version of Phrank that we next use to both discover patient cohorts and prioritize candidate disease genes without needing to share sensitive patient genotype and phenotype data.

### Secure multiparty computation

We construct our privacy-preserving protocols using techniques from modern cryptography, and specifically, secure multiparty computation. To illustrate the concept of secure multiparty computation, consider the following simple scenario: Patient 1 holds a secret number *X*, and Patient 2 holds a secret number *Y*. Patients 1 and 2 would like a test provider to compute the value *X* + *Y* for them. Of course, they can both simply give *X* and *Y* to the test provider who would then compute and reveal the sum *X* + *Y*. But in this model, the two patients have to *fully trust* the test provider, since the test provider learns both *X* and *Y* (Figure 1A). The objective in secure multiparty computation is to develop a protocol that allows the provider to provide Patients 1 and 2 the same service while protecting the secrecy of *X* and *Y* (namely, the provider should only be able to learn the output *X* + *Y*, but not *X* or *Y* themselves).

**Figure 1.**
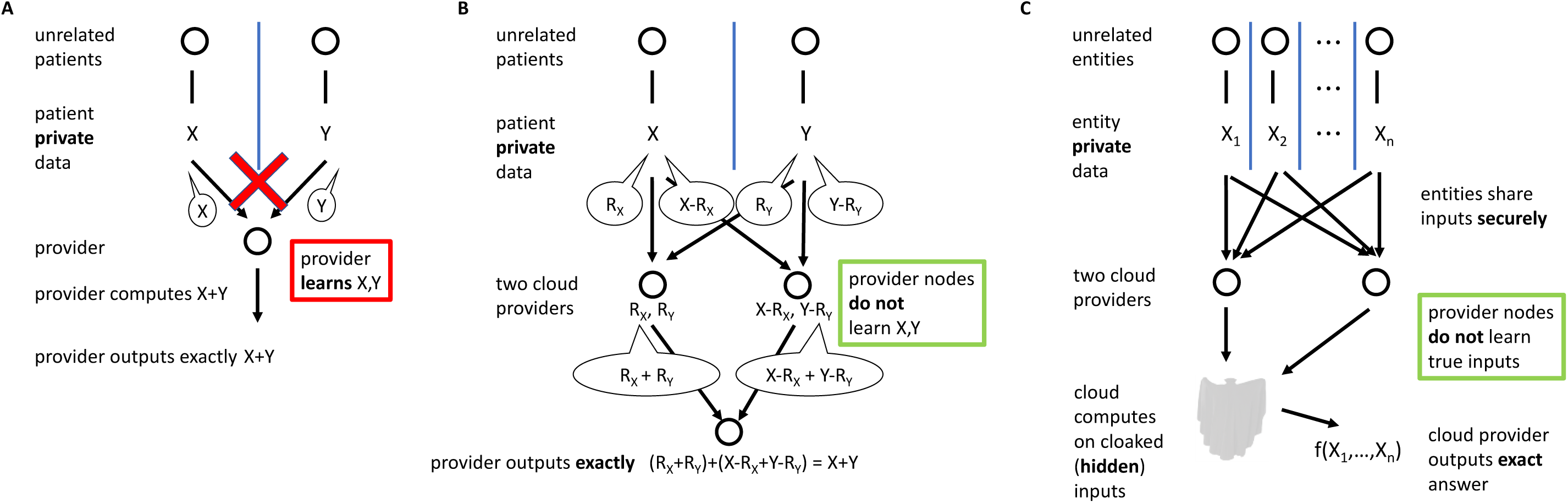
Secure multiparty computation with 2 clouds. In our simplified scenario, two patients rely on a cloud provider to perform some computation for them. (A) In existing protocols, outsourcing the computation to the cloud requires that the patients allow the provider to learn their private data. (B) Using two non-colluding clouds, patients can enjoy the same service without needing to share their private data. Because R_x_ and R_y_ are random numbers (see text), the provider can now exactly compute X + Y while learning nothing about X or Y themselves. (C) We extend this strategy here to more complex functions including any combination of addition and multiplication operations over numbers, vectors or matrices.

One way to achieve the above goal is to split the provider into two *non-colluding* test providers (assumed to be “honest-but-curious”—namely, they follow the protocol as described), which then jointly compute *X* + *Y* while individually learning nothing at all about *X* or *Y*. This privacy-preserving operation can be achieved using a notion called “secret sharing” (Figure 1B). In our simple example, Patient 1 picks a random number *R*_*X*_. She sends *R*_*X*_ to the first test provider service (Node 1), and she sends *X* − *R*_*X*_ to the second test provider (Node 2). Since *R*_*X*_ and *X* − *R*_*X*_ are both random numbers, they perfectly hide *X* from Node 1 and Node 2. Patient 2 does the same. He chooses a random number *R*_*Y*_ and sends *R*_*Y*_ to Node 1 and *Y* − *R*_*Y*_ to Node 2. Node 1 then computes *R*_*X*_ + *R*_*Y*_ and reveals only this number to Node 2. Similarly, Node 2 computes (*X* − *R*_*X*_) + (*Y* − *R*_*Y*_) and reveals only this number to Node 1. Nodes 1 and 2 now sum the two numbers they have just shared. The sum is exactly *X* + *Y*, but neither node has learned anything about either *X* or *Y* (see Figure 1B).

These ideas can be extended to support more complex computations involving any number of patients (Figure 1C), who first securely share some information with the two test provider services (Nodes 1 and 2). The two services securely compute on the shared inputs, and as long as they do not collude, the only information that is revealed to either of them is the outcome of the computation and nothing else. To reduce the cost of our protocols, we work in the “preprocessing” model(*14*–*17*) where we assume that prior to the computation between the two test providers, a third party called the “dealer” performs some precomputation and gives the output of the precomputation to the two test providers for use in the protocol. This precomputation does not depend on any participant’s secret inputs, and can be performed at any time. The dealer could be implemented by a third cloud provider or by patients (smartphones) participating in the protocol. This is the basic building block for the privacy-preserving protocols we develop in this work. We refer the reader to Methods for a rigorous discussion.

### Secure multiparty cohort discovery

Given a large number of undiagnosed patients, the goal of cohort discovery is to identify one or more small subsets of patients who share very similar phenotypes (and thus are likely to arise from a common mutated gene). The subsequent sequencing of these small cohorts in search of a common mutated gene has already led to the discovery and diagnosis of hundreds if not thousands of monogenic diseases.

Phrank enables an efficient approach for cohort discovery. Namely, we begin by computing the Phrank similarity score between (the sets of phenotypes of) all pairs of patients. Then, we identify small groups of patients who have high Phrank similarity scores with other members within the group, but low similarity scores with the overall large heterogeneous set of patients (Figure 2A). We develop a privacy-preserving way of implementing this cohort discovery algorithm that does *not* require patients to share their set of phenotypes with other patients or with the test administrator (Figure 2B). Specifically, we first express the Phrank computation(*13*) and the necessary clustering as a sequence of additions, multiplications, and Boolean operations. We then apply the “ABY approach”(*18*) to implement this protocol: we use the method from Figure 1 for secure addition, a method called Beaver triples(*14*) for secure multiplication, and Yao’s garbled circuits(*19*) for secure Boolean operations (see Methods and Supplementary Figure 1 for a complete description).

**Figure 2.**
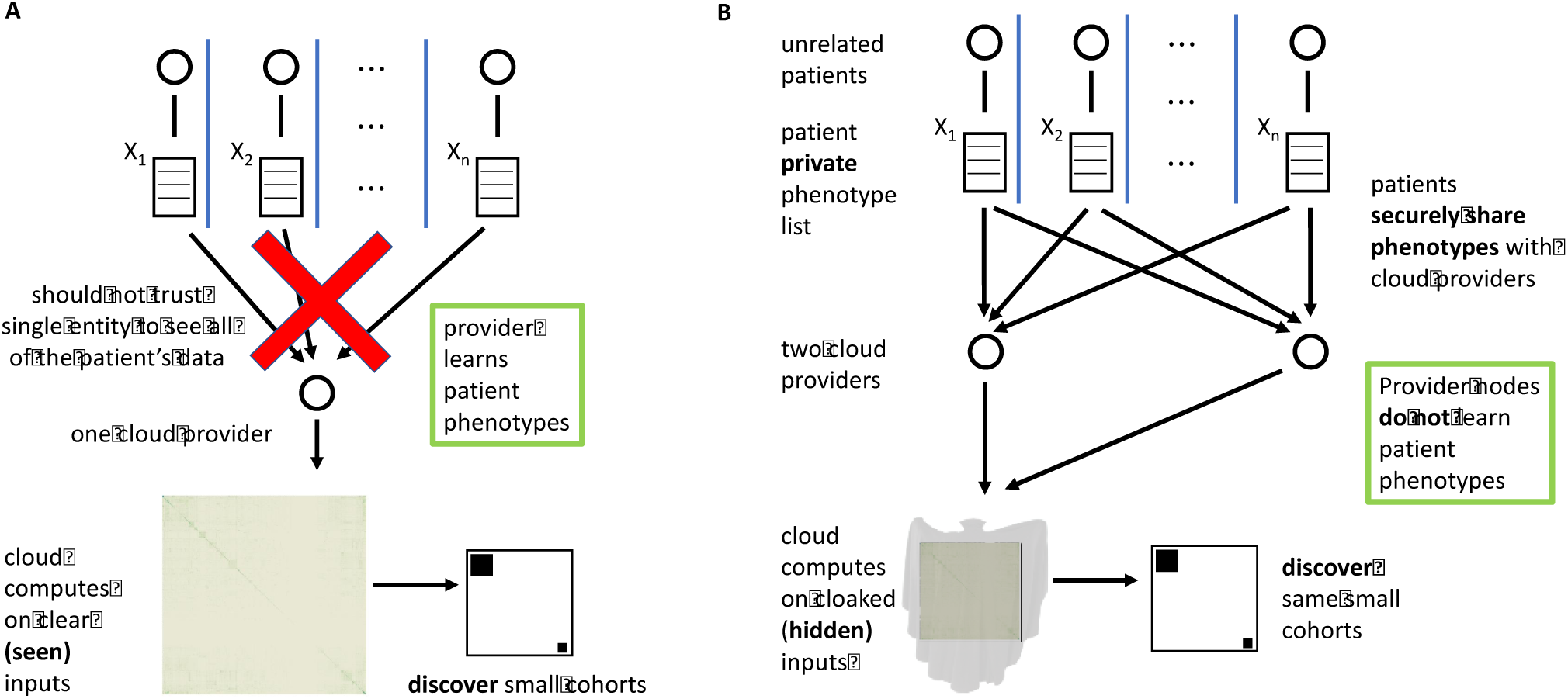
Scenario 1: Phenotypic cohort discovery without sharing any phenotypes. (A) Today, undiagnosed patients must share their phenotypes with one another and with test providers in hope of matching into a small cohort that can in turn be leveraged to identify the causative gene for their condition. (B) As in Figure 1, using 2 non-colluding cloud servers, patients can enjoy the same service without having to share their private data. In our scenario, undiagnosed patients can participate in cohort discovery multiple times, without revealing *anything* about themselves until they are matched (see Supplementary Figure 1 for technical details and Supplementary Figure 2 for real patient results).

We applied this technique to discover disease cohorts from a set of 1067 real patients with various Mendelian disorders, including 9 patients with Distal Arthrogryposis (DA) and 9 with Nager Syndrome (NS). The remaining 1049 patients were obtained from a large cohort of patients with likely genetic disorders (see Methods). Each of the 1067 patients was associated with a list of on average 4 clinician-noted HPO phenotypes, which was used to compute the pairwise phenotypic similarity between all patients. The maximum phenotypic similarity score between any two patients was 127.29, the average was 1.64, and the minimum was 0. The secure cohort building operation successfully identified 2 distinct cohorts among these 1067 patients (Supplementary Figure 2), where 8 of the 9 patients with Distal Arthrogryposis were grouped together and 9 of the 9 patients diagnosed with Nager Syndrome were grouped together. None of the 1049 patients with other diseases were incorrectly grouped into either cluster and no other cluster was incorrectly construed.

Under our experimental setup, placing Node 1 & 2 nearly 3,000 miles apart (one on each coast of the U.S), evaluating the secure patient cohort building operation on a group of 1067 real patients completed in just over 10 seconds and required 60 MB of online communication and 634 MB of precomputed values (see Table 1). Since the cohort building operation requires computing the phenotypic similarity between all pairs of patients, the bandwidth, size of the precomputed values, and the protocol execution time grow quadratically with the number of patients (see Supplementary Figure 3).

**Table 1.** Keeping patient phenotypes and genotypes private while seeking disease diagnoses. In Scenario 1, we provide as input the phenotypes of over 1,000 real patients, and successfully identify among them two small groups of patients with similar phenotypes. Our protocol does not require any patient to share their phenotypes with anyone else (Figure 2). In Scenario 2, a commercial provider accelerates the diagnosis of *n* = 169 patients without needing to see any patient phenotypes or genotypes (Figure 3). In this case, the patients do not learn much more about the commercial provider’s secret knowledgebase or exact approach, bar a small number of candidate genes that plausibly explain the patient’s condition. All computations are performed over real patient data.

|  | Relevant Information for Each Operation |  |  |  |  |  |  | Running Time Measurements |  |
| --- | --- | --- | --- | --- | --- | --- | --- | --- | --- |
| Scenario 1:<br>Patient cohort discovery | Diseases | Individuals in identified cohort | Individuals with Disease | Total Patients in Analysis | Avg. Similarity between Patients with Same Disease | Max/Avg./Min Similarity Across All Patients |  | Bandwidth (Megabytes) | End-to-End (sec) |
|  | Nager Syndrome | 8 | 9 | 1067 | 56.2 | 127.29/1.64/0 |  | 606 | 121 |
|  | Distal Arthrogryposis | 9 | 9 |  | 99.8 |  |  |  |  |
| Scenario 2:<br>Gene prioritization ($n = 169$ patients) | Total # of Known Protein-Coding Genes in Genome | Total # of known disease genes | Median # of Candidate Causative Genes per Patient (private) | Total # of HPO Phenotypes | Median # of Phenotypes per Patient | % of patients with causative genes in top 10 | Median Similarity Score between Patient and Causative Gene | Bandwidth (Megabytes) | End-to-End (sec) |
|  | 20,663 | 3,406 | 121 | 11,532 | 7 | 88% | 22.3 | 429 | 84 |

### Secure gene prioritization

Genome sequencing of an individual with a monogenic disease can yield hundreds of different candidate genes with rare functional mutations(*4, 5, 20*). With 60 million patients to be sequenced in the next 5 years(*21*), automated methods must be developed to assist the clinician in efficiently sifting through the candidate gene list in search of a possible diagnosis.

We have previously shown Phrank’s utility in this context. Namely, given a patient with a set of candidate genes and observed phenotypes, and given a knowledgebase consisting of all candidate disease genes and the set of phenotypes associated with each one, Phrank sorts the patient genes in descending order for their ability to explain the patient’s phenotypes with a very high likelihood of finding the patient’s causative gene at or near the top of this list(*13*). As described, this test asks the patient to reveal all of the genes where they have potentially causative variants as well as their full set of phenotypes to the test provider. Conversely, having the provider simply hand their full knowledgebase containing thousands of carefully-curated gene-phenotype associations to the patient would jeopardize their advantage over competing services (Figure 3A). Here, we use similar techniques as above to build a privacy-preserving framework for secure gene prioritization (Figure 3B). In this scenario the test provider does not learn *anything* about the patient, and the patient only learns the identities of the top few candidate genes that are most correlated with her particular phenotypes, and nothing more about the test provider’s knowledgebase or exact ranking method (see Methods, Supplementary Figure 4).

**Figure 3.**
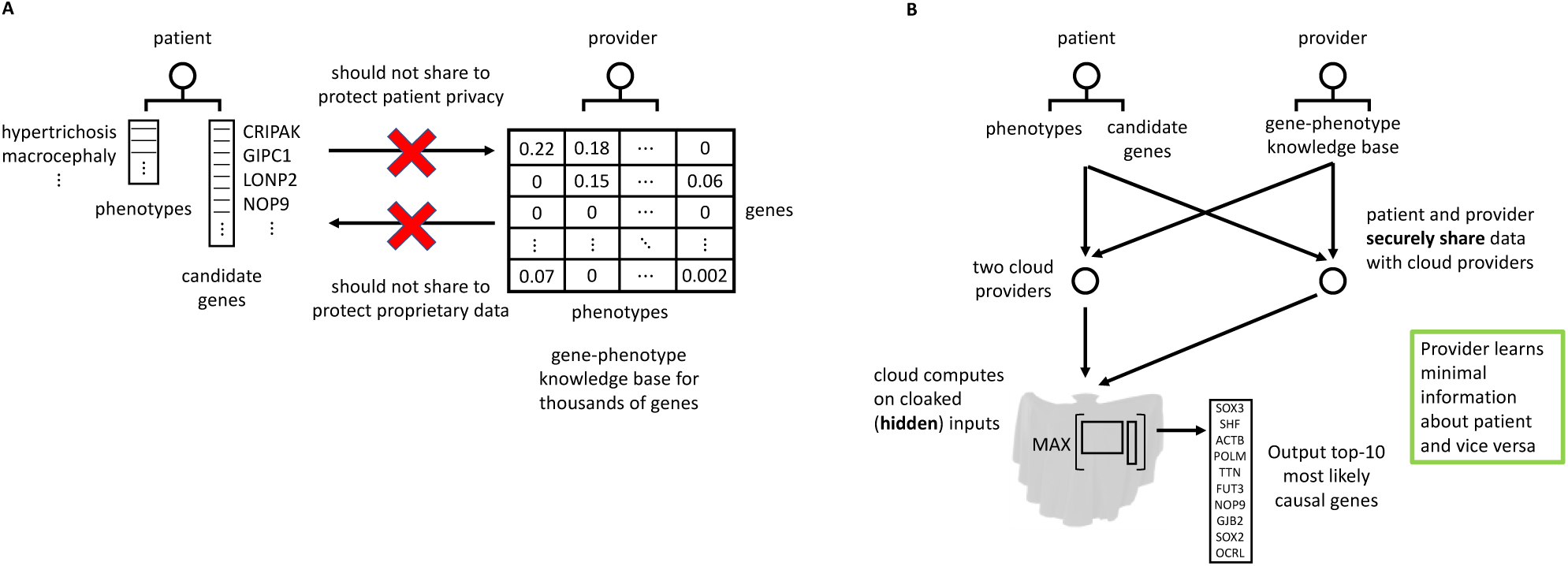
Scenario 2: Accelerating patient diagnosis while protecting both patient and commercial provider. Mendelian patients present hundreds of candidate genes, but only one of them may provide a diagnosis. Commercial providers compete over their ability to discover this causal gene. (A) Patient wants the provider to rank their candidate gene list, but they don’t want to reveal their private data. Neither does the provider want to entrust all their data and methodology to any single client. (B) Using our 2 cloud model, the patient protects all their sensitive data while the provider reveals only a sliver of theirs (see Supplementary Figure 4).

We ran our secure gene prioritization protocol over 169 real patients with different Mendelian disorders ranging from Albinism to Weaver Syndrome. The patients were associated with an average of 7 clinician-noted HPO phenotypes and typically presented between 100-150 candidate genes containing rare, potentially causative, variants. The secure computation outputted the top 10 of over 3400 disease genes that were most likely to explain each patient’s phenotypes. In nearly 90% of the cases (150 of the 169 cases), the causative gene was found in this 10-15X shorter list of genes. Depending on the precise privacy and accuracy requirements, the filtering steps for controlling the number of genes in the input of the secure gene prioritization protocol can be adjusted.

Our secure gene prioritization protocol completed in 84 seconds and required 429 MB of communication (see Table 1). The bandwidth, size of the precomputed values, and the protocol execution time all scale linearly with the size of the test provider’s knowledgebase (see Supplementary Figure 3).

## Discussion

Genomic data presents each family a ‘serve or protect’ dilemma. On the one hand, sharing the family’s genetic information with clinicians and researchers can advance genomic medicine and enable better treatment plans for the afflicted. On the other hand, the sensitivity of that same data requires prudence and care in handling so as to protect the patient family from genomic discrimination or exploitation. Our previous works have shown how to leverage secure multiparty computation techniques to identify disease-causing(*9*) or disease-associated(*22*) genes and obtain a definitive genetic diagnosis(*9*) without revealing nearly any patient genetic information. These works, however, implicitly assume that patient cohorts have already been assembled and that computation need only be done on the patients’ genotype data. In this work, we extend these secure genome techniques to enable secure computation over patients’ phenotypes as well. Combined, our protocols bring us closer to a secure *end-to-end* disease diagnosis pipeline capable of building disease cohorts, identifying their shared causal gene, and diagnosing subsequent patients(*9*), all while respecting the privacy of patients’ genetic and genomic data.

In combination with apps that store a patient’s medical record, and automated tools that extract HPO terms from medical notes (such as our ClinPhen(*23*)), one can envision a near future where a patient can decide to participate in a secure cohort discovery with a simple push of a smartphone button. As 70% of sequenced patients with severe genetic diseases remain undiagnosed(*10, 11*), and especially since 300 novel disease genes are discovered each year(*2*) (predominantly through cohort building), this is a highly desirable future where the latest technology both serves and protects us. Fierce competition between different commercial entities offering monogenic patient diagnosis make our secure gene prioritization protocol highly desirable to patients and commercial providers alike. The 30% of sequenced patients who can already receive a diagnosis will do so in an accelerated fashion, and without needing to share any private genomic or phenotypic data. Commercial providers in turn can amass knowledge in the form of better annotations, HPO graph refinements, and even enhancements to Phrank while only revealing a tiny sliver of vital information to each patient.

In both scenarios, virtually nothing is discovered about the patients, especially in cases where the computation cannot currently aid the patients directly. This allows patients to participate in multiple cohort building efforts, until a cohort is found that can aid them too. It should also allow different healthcare systems to join forces in building small patient cohorts, without ever sharing any actual patient data. And it allows patients in our second scenario to get reanalyzed periodically by the same, or alternatively, a different, commercial entity to facilitate a diagnosis when one ultimately becomes available.

The methods we describe in this work enable us to significantly enhance the disease diagnosis pipeline while protecting the privacy of all parties involved. The core primitive that underlies our protocols is the Phrank phenotype set-similarity metric, and the computational overhead invested to make the computation secure is very feasible in terms of computation time, bandwidth, and the amount of precomputation. It is straightforward to further generalize Phrank to support more functionalities and computations over phenotypes. For example, using the AMELIE PubMed article/disease gene/phenotypes database(*24*), publications can also be securely prioritized to reveal only those that best explain a patient’s disease.

This work introduces the first framework for computing meaningful functions over Mendelian phenotypes and genotypes without requiring patients to reveal their phenotype or genotype information. These protocols enable accurate disease diagnosis while revealing minimal patient information, a large step towards enabling secure personalized genome analysis for all.

## Online Methods

### Human Phenotype Ontology

Phenotypes are represented using the Human Phenotype Ontology(*12*) (HPO) Build 127. HPO phenotypes are arranged in a directed acyclic graph (DAG), where the graph structure encodes the parent-child relationship between the different phenotypes. The DAG is organized such that if a patient or gene is associated with a phenotype *φ*, the patient or gene is also associated with all ancestors of *φ* up to the root phenotype node “phenotypic abnormality” (HP:0000118).

### Patient phenotype data

Patient phenotypic data were combined from 4 sources. Phenotypes for 1049 patients were extracted from patient records found in the Stanford STARR database(*23*). 169 patients from the Deciphering Developmental Disorders(*25*) (DDD) cohort came associated with a list of HPO phenotypes. We manually extracted HPO phenotypes for 9 patients identified with Nager Syndrome from Table 1 in Bernier et al(*26*) and for 9 Distal Arthrogryposis patients from Table 1 in McMillin et al(*27*).

### Patient genotype data

Variant Call Format (VCF) files of patients submitted to the Deciphering Developmental Disorders (DDD) project were downloaded from European Genome-Phenome Archive (EGA) with accession numbers EGAD00001001848, EGAD00001001977, EGAD00001002748, EGAD00001001355, EGAD00001001413 and EGAD00001001114. All patients with a single-gene diagnosis not due to a structural variant (as specified by the patient’s VCF) and for which the causative gene was not a novel discovery of the DDD project were selected. From any diagnosed set of siblings, a single diagnosed sibling was selected at random resulting in 169 diagnosed DDD patients as shown previously(*13*).

### Variant annotation

ANNOVAR v527 was used to annotate patient variants with predicted effect (below) on protein-coding genes using gene isoforms from ENSEMBL gene set version 75 for the hg19/GRCh37 assembly of the human genome. We included all gene isoforms where the transcript start and end were marked as complete and where the coding span was a multiple of three. All ExAC v and 1000 genomes phase 3 subpopulations were used to annotate variants with allele frequency information.

### Variant filtering

Only variants with an exonic nonsynonymous SNV, exonic stopgain, exonic frameshift, core splicing, exonic nonframeshift or exonic stoploss predicted semantic effect on a protein-coding gene isoform were considered for further analysis. All patient variants with an allele frequency over 0.5% in any subpopulation of the 1000 Genomes Project or in Exome Aggregation Consortium (ExAC) were also marked as likely benign. In case a gene contained only a single heterozygous variant, the variant was marked as likely benign if it occurred at an allele frequency over 0.1% in any subpopulation of the 1000 Genomes Project or ExAC. All variants marked as likely benign were filtered out of the candidate list of variants. All remaining variants made up the list of candidate causative variants for each patient.

### Additive secret sharing

A two-party secret-sharing protocol allows a user to distribute “shares” of a value *x* to two servers such that each server individually has no information about *x*. Collectively, the two servers can combine their shares to reconstruct the secret *x*. In this work, we use an additive secret-sharing scheme. To secret share *x* across two parties, the user chooses a random value *r* between 0 and *p* – 1 (where *p* is a fixed integer). In our protocols, we use *p* = 2^16^. The user then gives *r* to one party and *x* – *r* (mod *p*) to the other party. Since *r* is uniformly random, neither party learns anything about *x* given just their share of the input. However, by adding their shares together, the parties can recover the secret *x* = *r* + (*x* – *r*) (mod *p*). We write [*x*] = ⟨[*x*]_1_, [*x*]_2_⟩ to denote a secret-sharing of *x*, where [*x*]_1_ denotes the share held by the first party (e.g. *r*) and [*x*]_2_ denotes the share held by the second party (e.g. *x* − *r* (mod *p*)). The invariant we maintain is that [*x*]_1_ + [*x*]_2_ = *x* (mod *p*).

### Computing on secret-shared values

#### Addition on secret-shared values

We now describe how two parties can jointly compute on secret-shared values without learning anything about the underlying shared values. Given shares of [*x*] = ⟨[*x*]_1_, [*x*]_2_⟩ and [*y*] = ⟨[*y*]_1_, [*y*]_2_⟩, the first party will have [*x*]_1_, [*y*]_1_ and the second party will have [*x*]_2_, [*y*]_2_. First, we remark that the shares are *additive* meaning that a party who has shares of *x* and *y* can just add their shares together to obtain a share of the sum *x* + *y*, or [*x* + *y*] = ⟨[*x*]_1_ + [*y*]_1_, [*x*]_2_ + [*y*]_2_⟩. Notably, this operation does *not* require any communication between the two parties. Similarly, parties can scale their shared values by a fixed constant by scaling their local shares. Namely, for a constant *k*, we can write [*kx*] = ⟨*k*[*x*]_1_, *k*[*x*]_2_⟩. Finally, to add a constant *y* to a secret-shared value [*x*], the two parties just need to compute [*x*] + *y* = ⟨[*x*]_1_ + *y*, [*x*]_2_⟩. In summary, these operations enable two parties to *locally* compute any *affine* function (i.e., linear functions of the form *ax* + *b*, where *a* and *b* are fixed constants) on their secret-shared values.

#### Multiplication on secret-shared values

Computing a product of two secret-shared values is more complex. Here, we describe an elegant technique due to Beaver(*14*) that we use in this work. Beaver’s multiplication protocol is a core ingredient in secret-sharing-based multi-party-computation protocols(*15*). First, we assume that the parties have a secret-sharing of a *random* product: namely, a triple ([*a*], [*b*], [*c*]), where *a* and *b* are *random* values (that do not depend on any party’s secret information), and *c* = *ab* (mod *p*). We refer to ([*a*], [*b*], [*c*]) as a “Beaver multiplication triple.” Beaver’s main insight is that we can leverage a secret-sharing of a *random* product to compute a secret-sharing of an arbitrary product. We describe the main protocol below:

1. Suppose that the two parties possess a secret-sharing of two values [*x*] and [*y*], and their goal is to compute a secret-sharing of the product [*xy*]. Moreover, the two parties have shares of the multiplication triple ([*a*], [*b*], [*c*]) where *c* = *ab*. Note that the two parties only have shares of the *a, b*, and *c*, and importantly, do not know the actual values of *a, b*, and *c*.
2. Each party computes and publishes their shares of *x* − *a* (mod *p*) and *y* − *b* (mod *p*). Specifically, the first party sends [*x*]_1_ − [*a*]_1_ and [*y*]_1_ − [*b*]_1_ to the second party, and the second party sends [*x*]_2_ − [*a*]_2_ and [*y*]_2_ − [*b*]_2_ to the first party. At the end of this step, both parties have the values *x* − *a* (mod *p*) and *y* − *b* (mod *p*).
3. The parties then locally compute the following *affine* relation:

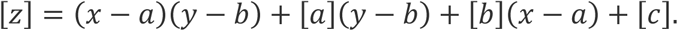 Specifically, the two parties compute the following:

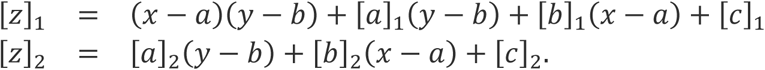

We argue that *z* is a sharing of the product *xy*. Appealing to linearity of the underlying operations and using the fact that *c* = *ab*, we show that *z* is a secret-sharing of the product *xy*:

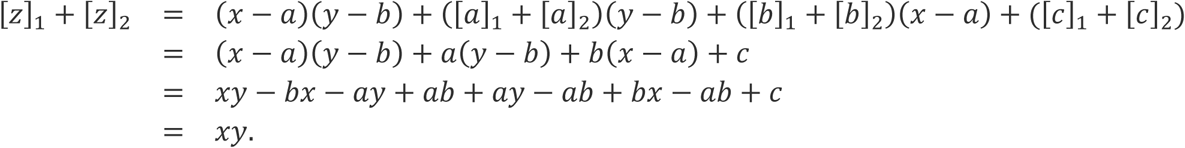

As long as the values *a, b*, and *c* in the multiplication triple are unknown to the two parties, then the values *x* − *a* and *y* − *b* completely hide *x* and *y* (in an information-theoretic sense). To summarize, Beaver’s protocol enables two (or more) parties to jointly compute a product of two secret-shared values with one round of communication (where each party broadcasts their shares of the blinded values to the other parties). With Beaver’s protocol, the problem of multiplying secret-shared values essentially reduces to that of generating the (correlated) Beaver multiplication triples. Note that the Beaver triples are completely *independent* of all of the parties’ inputs to the computation, and thus, can be generated in a separate *offline* phase prior to the main protocol execution. We discuss this more below (see “Using preprocessing for better online efficiency”).

#### Generalizing Beaver multiplication triples to matrix-vector operations

In the above, we described a protocol for computing on secret-shared values (specifically, field elements). A significant component of our protocols is evaluating matrix-matrix and matrix-vector products. While we can perform the matrix operations using the elementary operations over the underlying elements in the matrices, this incurs unnecessary overhead. To improve performance, we note that Beaver’s multiplication protocol directly generalizes to computing products of secret-shared matrices (in fact, it extends to computing products over any ring), provided that the computing parties have a secret-sharing of a *random* matrix product (of the same dimensions). Specifically, for matrices ***X*** and ***Y***, we write [***X***] and [***Y***] to denote a secret-sharing of ***X*** and ***Y***, respectively (this is just a component-wise secret-sharing of the entries in the matrix). Addition and scalar multiplication on secret-shared matrices and vectors can be done exactly as before. For multiplication, we use the direct generalization of Beaver’s protocol. In particular, we assume that the computing parties have a secret-sharing of a matrix [***A***], and vectors [***B***] and [***C***] where ***C*** = ***AB***. To compute shares of the product [***XY***] given shares of matrices [***X***] and [***Y***], the two parties first reveal ***X*** − ***A*** and ***Y*** − ***B***. Then, they compute the following affine relation on their shares:

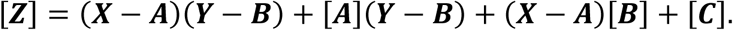

Using the same analysis as above, we have that [***Z***] = [***XY***], as desired. The total communication to perform a matrix vector multiplication in this way is proportional to the dimensions of ***X*** and ***Y***, which can be significantly smaller than the number of elementary multiplications that need to be done to compute the product ***XY***. In particular, if ***X*** is *m*-by-*k* and ***Y*** is *k*-by-*n*, then the total communication is *k*(*m* + *n*), whereas using Beaver multiplication triples to perform each of the elementary multiplications requires communication 2. *kmn*. This optimization (along with further generalizations) was also leveraged to improve the efficiency of the privacy-preserving GWAS protocol by Cho et al(*22*).

#### Yao’s garbled circuits for evaluating Boolean circuits

Beaver’s protocol provides an efficient method for evaluating arithmetic operations (addition and multiplication) on secret-shared values. However, they are less suitable for other types of operations, such as comparisons or computing the maximum of a collection of values. While it is possible to express these operations in terms of additions and multiplications, doing so incurs considerable overhead (in terms of both computation as well as communication). A more efficient alternative for secure evaluation of comparisons and argmax operations is to use Yao’s protocol(*19, 28*), which is a general-purpose protocol for secure two-party computation. At a high level, suppose Alice and Bob each have an input *x* and *y*, and they want to compute some joint function *f* on their shared inputs. At the end of the computation, both Alice and Bob should learn *f*(*x, y*) but nothing else about the other party’s input (other than what is revealed by *f*(*x, y*)). In Yao’s protocol, the function *f* is modeled as a *Boolean* circuit, and the inputs are binary-valued. In contrast, in the case of Beaver’s protocol, the underlying function is modeled as an arithmetic circuit (over a large finite field). Because comparisons are more naturally expressed as a Boolean circuit, Yao’s protocol is more suitable for performing the comparison and argmax operations.

### The ABY approach: combining Boolean and arithmetic circuits

In this work, the computation we are interested in consists of two main ingredients: evaluating a matrix-vector (or matrix-matrix) product, and then applying some post-processing operations to the results (e.g., computing the top *k* elements in the resulting vector or filtering out values that fall below a certain threshold). Since matrix-vector products are arithmetic computations, they are most well-suited for secret-sharing-based secure multiparty computation. On the other hand, computing comparisons and thresholds are more easily expressed as Boolean circuits (that operate over the binary representation of the inputs), and thus, are more amenable for Yao’s protocol. The ABY approach(*18*) combines the best of both worlds and provides a general paradigm for integrating secret-sharing multi-party computation with Yao-based multi-party computation. In this work, we rely on the following elementary building blocks from Demmler et al(*18*).

- **Evaluating a garbled circuit on secret-shared values.** First, we describe how to evaluate a Boolean function on secret-shared values. Let *f* be the Boolean function (e.g., this could be an argmax function or a threshold function) we want to evaluate, and suppose the inputs [***v***] = ([***v***]_1_, [***v***]_2_) to *f* are secret-shared across two-parties. To use Yao’s protocol to evaluate the function *f* on the secret-shared values, we first define the function

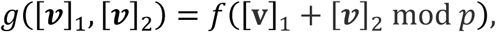

and then apply Yao’s garbled circuit protocol to evaluate the function *g*. In this case, the input of each party is the (binary) representation of their share of the input. At the end of the protocol execution, the parties learn the output

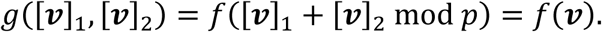
- **Secret-sharing the output of a garbled circuit.** In some cases, the output of the Boolean circuit is not the final output of the computation, and we need to perform additional arithmetic operations on the result. In this case, the outputs of the Boolean circuit should be a *secret sharing* of the output rather than the output. We use this operation to implement the PRIORITIZATION functionality (described further below). Suppose we want to compute a Boolean function *f* on an input ***v***, and we want the output ***z*** = *f*(***v***) to be secret-shared across the two parties. To support this, we define a new function *g* that takes in the input ***v*** as well as a share [***z***]_2_ as follows:

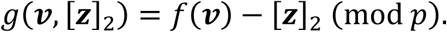

The input [***z***]_2_ belongs to the second party, and is sampled uniformly at random (by the second party). By construction, the output of *g* is a vector [***z***]_1_ where [***z***]_1_ + [***z***]_2_ = ***z*** (mod *p*) and moreover, [***z***]_2_ is sampled uniformly at random. Thus, the pair ([***z***]_1_, [***z***]_2_) is a secret-sharing of ***z***. At the end of the computation, the first party obtains the share [***z***]_1_ while the second party holds the share [***z***]_2_ (unknown to the first party).

### Using preprocessing for better online efficiency

We can often reduce the cost of a secure multiparty computation (MPC) protocol by working in the MPC with preprocessing model(*14*–*17*). In this model, there is an independent semi-honest party (called the “dealer”) that generates some *input-independent* values for the computing parties. In other words, the protocol can be decomposed into an initial “offline phase” and an “online phase” which operate as follows:

- **Offline phase.** During the offline phase of the protocol, the dealer precomputes the input-independent values and distributes them to the computing parties. At the end of the offline phase, the dealer can go offline and it does not have to be around for the online phase of the computation.
- **Online phase.** During the online phase of the computation, the computing parties obtain the secret-shared inputs from all the protocol participants (e.g., the patients or the doctors). They then run the protocol over the secret-shared data. At the end of the computation, the computing parties publish their shares of the output, which allows the clients to learn the output of the computation. During this phase, the computing parties can make use of the precomputed values they received during the offline phase of the protocol.

We now briefly describe how we take advantage of preprocessing in our protocols:

- **Beaver’s multiplication protocol.** Recall from our above description that Beaver’s protocol for multiplying secret-shared values assumes the parties have a secret-sharing of a random multiplication triple (which can be generated independently of all of the inputs to the computation). These multiplication triples would be generated by an independent semi-honest party and distributed to the computing parties prior to the main protocol execution.
- **Yao’s garbled circuit protocol:** Working in the preprocessing model also enables a more efficient implementation of Yao’s garbled circuit protocol (which we use for securely evaluating Boolean circuits). Notably, since the garbled circuits used in our protocol are fixed (and input-independent), they can be precomputed prior to the computation. This means that in the online phase of the protocol, the two computing parties do not have to communicate the description of a large garbled circuit, which considerably reduces the online communication, and correspondingly, the end-to-end online protocol execution time. Finally, we can also reduce the online cost of the oblivious transfers (OTs)(*29, 30*) used to implement Yao’s garbled circuit protocol by precomputing OT correlations(*31*). With precomputed OT correlations, each of the 1-out-of-2 OTs on the input wire encodings to the garbled circuit (128-bits each) can be implemented by communicating 257 bits (1 bit sent from the evaluator to the garbler and 256 bits sent from the garbler to the evaluator). No cryptographic operations are needed. Using precomputed OT correlations to implement the OTs reduces both the communication and the computational cost of implementing the OTs.

In our experiments, we work in the preprocessing model and assume that there is a dealer that generates the Beaver multiplication triples for the arithmetic computations and the OT correlations as well as precomputed garbled circuits for the Boolean computations. In practice, this dealer could be implemented by a third independent cloud provider or by the patients (smartphones) who are contributing their data to the protocol execution. Finally, in our experiments, we measure the total online cost of the computation as well as the total size of the precomputed values that need to be distributed prior to the start of the online computation.

### Implementing the application-specific wrappers

We now describe how we implement our application-specific wrappers (patient COHORT DISCOVERY and GENE PRIORITIZATION) as a combination of arithmetic and Boolean circuit operations.

### Representing phenotypic and genotypic data as vectors

Phenotypes are represented using 11,532 distinct terms in the Human Phenotype Ontology (HPO) terms (see above). For each set of HPO phenotype terms **Φ**, we define two 11,532-dimensional vectors, ***v***_Φ_ and ***w***_Φ_, where each vector component corresponds to an HPO term. An *indicator phenotype vector* ***v***_Φ_ is a vector where the component corresponding to an HPO term *φ* is 1 if *φ* ∈ Φ, and 0 otherwise. A *weighted phenotype vector* ***w***_Φ_ is a vector where the component corresponding to an HPO term *φ* is its “weight” *w*_*φ*_ if *φ* ∈ Φ, and 0 otherwise (see Methods). The weight of a phenotype, *w*_*φ*_, is defined to be the marginal information content of the phenotype node (as described in Phrank)(*13*).

There are 20,663 protein-coding genes in the Ensembl 75 gene set. Where gene-level genomic data is required for the computation, we create a 20,663-dimensional indicator genotype vector where each vector component corresponds to a gene. The components corresponding to the genes harboring a rare functional variant are set to 1 and all other components are set to 0 (see below).

#### Scenario 1: Secure patient COHORT DISCOVERY

Recall that the COHORT DISCOVERY functionality takes as input a set of phenotypes from multiple patients, computes the pairwise phenotype similarity score between each pair of patients, and then identifies small clusters of patients. Suppose there are a total of *n* patients. Let ***v***_1_, …, ***v***_*n*_ be the indicator phenotype vector (i.e., a 0/1 vector) that models the patient’s phenotypes (as above). Let ***w***_1_, … ***w***_*n*_ be the weighted phenotype vector of for each patient (an entry in ***w***_*i*_ is 0 if the patient does not exhibit the particular phenotype, and is equal to the weight of the phenotype if the patient does have the particular phenotype). We define the COHORT DISCOVERY functionality as follows:

1. Let ***V*** be the matrix whose columns consist of the vectors ***v***_1_, …, ***v***_*n*_ and ***W*** be the matrix whose columns consist of the vectors ***w***_1_, …, ***w***_*n*_.
2. Compute the pairwise-similarity matrix ***S*** = ***V***^*T*^***W*** consisting of the Phrank scores between each pair of patients. Note that ***S*** is an *n*-by-*n* matrix.
3. To identify clusters of similar patients, apply the following filtering operation to the entries of ***S***. First, let *s*_max_ be the maximum value among the entries in ***S***, and define the threshold *t* = *τ* · *s*_max_ for some parameter 0 < *τ* < 1. Let ***S***_thresh_ be the *n*-by-*n* matrix where (***S***_thresh_)_*i,j*_ = 1 if ***S***_*i,j*_ > *t* and (***S***_thresh_)_*i,j*_ = 0 otherwise. Namely, ***S***_thresh_ is an *n*-by-*n* indicator matrix whose 1-entries precisely correspond to the entries in ***S*** that exceed the threshold *t*.
4. Finally, to filter out small clusters, define the *n*-by-*n* indicator matrix ***S***_filtered_ where (***S***_filtered_)_*i,j*_ = 1 if (***S***_thresh_)_*i,j*_ = 1 and there are at least *ρ* non-zero entries in the *i*^th^ row *and* the *j*^th^ column of ***S***_thresh_. Otherwise (***S***_filtered_)_*i,j*_ = 0. Intuitively, this step filters out all pairs of patients that are not sufficiently similar to at least *ρ* other patients each. The output of the algorithm is the filtered matrix ***S***_filtered_.

In the above description, both thresholds *τ* and *ρ* are fixed parameters chosen based on the specifics of the particular scenario. In our empirical experiments, we use *τ* = 1/4 and *ρ* = 4.

We now describe our protocol for secure evaluation of the patient COHORT DISCOVERY functionality (see Figure 2). We work in the two-cloud model with preprocessing, where we assume that the online computation is performed between two non-colluding servers and that there is a third independent server (the “dealer”) that implements the offline precomputation and distributes the precomputed values to the two cloud providers prior to the start of the online computation.

At a high level, the online phase of our protocol works as follows. Each of the patients begins by secret-sharing their indicator phenotype vector and their weighted phenotype vector to the two non-colluding clouds servers. Next, the two clouds leverage Beaver’s multiplication protocol to compute a secret-sharing of the pairwise-similarity matrix ***S*** for the patients. Finally, the two parties apply Yao’s garbled circuits to the shares of ***S*** to perform the thresholding and filtering and obtain the final 0/1 matrix ***S***_filtered_. We describe our protocol formally below:

1. Each party secret shares its inputs ***v***_*i*_ and ***w***_*i*_ with the two cloud servers. At the end of this step, the two clouds have shares [***v***_1_], …, [***v***_*n*_] and [***w***_1_], …, [***w***_*n*_] of every party’s input. Equivalently, each cloud has a secret share of the matrices [***V***] and [***W***], where ***V*** is the matrix whose columns consist of the vectors ***v***_1_, …, ***v***_*n*_ and ***W*** is the matrix whose columns consist of the vectors ***w***_1_, …, ***w***_*n*_.
2. Using Beaver’s multiplication protocol, the two clouds compute a secret sharing of the product [***S***] = [***V***^*T*^]. [***W***].
3. Let *f*_filter_[*τ, ρ*] be the function that takes as input a pairwise-similarity matrix ***S*** and performs the thresholding and filtering procedure described above (using thresholds *τ* and *ρ*). Then, define the filtering procedure *g*_filter_[*τ, ρ*] that operates on secret-shared values as follows:

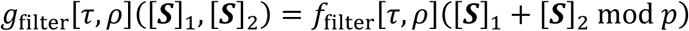 The two clouds use Yao’s garbled circuit protocol to jointly evaluate *g*_filter_[*τ, ρ*] where the first cloud provides its share [***S***]_1_ as its input and the second party provides its share [***S***]_2_ as its input.

In the offline phase of the protocol, the dealer generates the Beaver multiplication triples, OT correlations, and garbled circuits necessary to implement the above-described protocol and distributes the precomputed values to the two computing clouds for use in the online phase of the protocol. We now provide some additional details on how we implement (and optimize) the above protocol:

- Phenotype weights are typically represented as decimal values while our secure computation protocol computes over integer values (in fact, integers modulo *p*). While it is possible to implement floating-point operations as integer operations, this incurs significant overhead. Instead, in this work (and similar to many previous protocols for secure computation), we use a *fixed-point* representation. In fact, since our protocols only care about the ordering and (relative) magnitudes of the information-scores, we simply scale the phenotype weights by a large multiplicative factor (in our work, we use 2^9^ = 512), and then round the weights to the nearest integer. Finally, we choose the modulus *p* to be large enough so none of the arithmetic operations wrap around modulo *p*. Note that whenever we have to operate on secret-shared values in Yao’s protocol, we need to perform a modular reduction (mod *p*) within the garbled circuit to reconstruct the input. It is thus convenient to choose *p* to be a power of two (or close to a power of two) so that the modulus reduction can be efficiently represented as a *Boolean* circuit. In this work, we use *p* = 2^16^.
- Computing the threshold *t* = *τ* · *s*_max_ for an arbitrary threshold 0 < *τ* < 1 will require implementing a division operation within a Boolean circuit, which can be very expensive in general. However, if we choose the threshold *τ* to be an (inverse) power of two, then division reduces to performing a bit-shift (more precisely, we simply drop some of the least significant bits). This can be implemented with almost no overhead. For this reason, we use *τ* = 1/4 in this work. If we want to efficiently support arbitrary thresholds that are not close to a power of two, another option is to modify the garbled circuit to first compute and output the maximum entry *s*_max_. Then, the parties can compute the threshold *t* in the clear and finally, run the filtering protocol with the threshold given as an additional input to the computation. Note that this modified protocol would (only) reveal the value (but not the identity) of *s*_max_ to the computing parties.
- The rest of the threshold function corresponds to performing comparisons, maximums, and counts. All of these elementary operations can be implemented efficiently using building blocks from previous works(*32*).

#### Scenario 2: Secure PRIORITIZATION

The PRIORITIZATION functionality can be viewed as a two-party computation between a patient and a 3^rd^ party genome analysis provider. In this scenario, the patient wants to obtain a short list of genes from the genome analysis provider, sorted based on the likelihood that a particular gene causes their disease. Moreover, the patient would like to do so without revealing their genotypic or phenotypic information to the genome analysis provider. In today’s competitive commercial environment, the gene-phenotype mappings, HPO structure and even enhanced Phrank algorithms used by the genome analysis provider often contains proprietary information, and we desire a protocol that additionally protects the confidentiality of the provider’s data.

In this scenario, the patient holds an indicator phenotype vector ***v***_*p*_ (i.e., this is a 0/1 vector specifying the set of phenotypes a patient possesses) and an indicator genotype vector ***v***_*g*_ (i.e., this is a 0/1 vector specifying the set of genes containing rare functional variants that the patient possesses). The genome analysis provider holds a set of weights matching genes to specific phenotypes. We represent this as a matrix ***S***. The rows of ***S*** are associated with genes while the columns of ***S*** are associated with phenotypes. Essentially, we can view each row of ***S*** as the weighted phenotype vector associated with the corresponding gene in the provider’s gene-phenotype database.

In our setting, the phenotype vector has dimension 11,532 (corresponding to the number of HPO terms in build 127), while the genotype vector has dimension 20,663 (corresponding to the number of protein-coding genes in Ensembl build 75). The PRIORITIZATION functionality identifies the genes the patient possesses that are most correlated with her phenotypes. The components of the vector ***Sv***_*p*_ can be viewed as the Phrank score(*13*) between the patient’s phenotypes and the phenotypes associated with each gene in the provider’s gene-phenotype database.

The PRIORITIZATION functionality begins by computing the Phrank scores vector ***Sv***_*p*_. Next, we filter out all genes where the patient does not have a rare variant by computing the Hadamard product ***w*** = ***v***_*g*_ ∘ (***Sv***_*p*_). To recall, the Hadamard product ***u*** ∘ ***v*** between two vectors ***u*** and ***v*** is defined to be the component-wise product of ***u*** and ***v***. The vector ***w*** encodes the Phrank score for each gene in the patient’s genome where the patient possesses a rare variant. The PRIORITIZATION functionality outputs the top *k* genes (entries) in the vector ***w***, which corresponds to the genes with the highest Phrank scores with respect to the patient’s set of phenotypes. Below, we will write TOP-K to denote the function that takes as input a vector ***v*** and outputs a 0/1 vector (of the same dimension as ***v***) indicating the *k* largest entries in ***v***.

In many cases (including the scenarios we consider), the phenotype weights are available only for a subset of the genes (e.g., for only the known disease-causing genes). While this can be handled in the basic protocol described above by setting the rows of ***S*** to be all zeroes whenever a gene is not present, this incurs additional cost in the secure computation (since the parties still need to compute a full matrix-vector product ***Sv***_*p*_). A more efficient method is to define a “projection matrix” **Π** that maps a 0/1 vector over the full set of genes (20,663 genes) to a 0/1 vector over a reduced set of disease genes (e.g., 3,406 genes). By construction, each row of **Π** is a 0/1 vector with a single 1 in one position (corresponding to the gene that it is selecting for). Then, the number of rows in the matrix ***S*** is equal to the number of genes in the *reduced* set (rather than the *total* number of genes in the genome). Computing the PRIORITIZATION functionality corresponds to computing

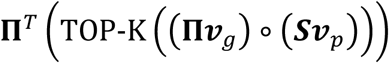

where ***S*** is the phenotype weights for the reduced set of genes. The matrix **Π**^*T*^ projects gene indices in the reduced set of genes to indices in the full set of genes. There are two main advantages to structuring the computation in this this way:

- All of the matrix-vector multiplications involving **Π** and **Π**^*T*^ are over binary matrices and vectors. Evaluating binary matrix-vector products requires less communication than evaluating matrix-vector products modulo *p* (namely, we can use a Beaver multiplication triple over bits rather than over values modulo *p*). Introducing the projection matrices allows us to perform a *smaller* matrix-vector multiplication modulo *p* (in exchange for performing two additional matrix-vector multiplications over binary inputs).
- The TOP-K computation only needs to be performed over the reduced set of genes. Since this is the bottleneck in the computation and in most cases, the subset of genes we are interested in is much smaller than the total number of genes, this yields a significant savings in both communication and computation.

We now describe how we securely evaluate the PRIORITIZATION functionality. There are several possible ways to perform the computation: either the patient can directly interact with the 3^rd^ party genome analysis provider, or it can secret-share its data to a cloud server and the genome analysis provider, and the resulting two-party computation occurs between the cloud server and the genome analysis provider. We take the latter approach here (see Figure 3), and in addition, we ask the client to play the role of the dealer in the offline phase of the protocol. Namely, the client (computer) generates the Beaver multiplication triples, the OT correlations, and the garbled circuit to be used in the online protocol execution.

1. At the beginning of the protocol, the two parties have secret shares of the patient’s indicator phenotype vector [***v***_*p*_], the patient’s indicator genotype vector [***v***_*g*_], the phenotype weights [***S***], and the projection matrix [**Π**].
2. Using Beaver’s multiplication protocol, the two parties compute a secret sharing of the vector [***z***] = ([**Π**] · [***v***_*g*_]) ∘ ([***S***]. [***v***_*p*_]).
3. The second party chooses a random 0/1 vector [***z*′**]_2_. This will correspond to its share of the output ***z*′** from the TOP-K computation. The two parties then use Yao’s garbled circuit protocol to evaluate the function

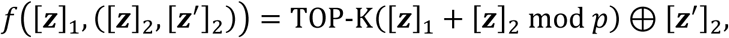

where *a* ⊕ *b* denote addition modulo 2. At the end of the protocol execution, the first party has [***z***′]_1_ = TOP-K([***z***]_1_ + [***z***]_2_ mod *p*) ⊕ [***z***′]_2_ while the second party has [***z***′]_2_. Equivalently, the two parties have a mod-2 secret-sharing of the vector ***z***′ = TOP-K(***z***). The basic Boolean operations needed to perform this computation correspond to performing additions modulo *p*, computing the max value in a vector, and comparisons between values. All of these operations can again be implemented using standard garbled circuit building blocks^24^.
4. To complete the protocol, the two parties use Beaver’s multiplication protocol to compute a secret-sharing of [**Π**^*T*^***z***′] = [**Π**^*T*^] · [***z***′]. The parties then broadcast their shares of the output. This suffices to reveal

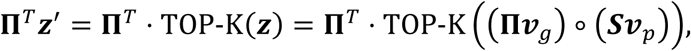

which is precisely the desired computation.

### Experimental setup

As discussed above, we implement our protocols in the two-cloud model, where the patients and data contributors first secret-share their data with two non-colluding cloud servers which perform the bulk of the computation. In addition, we assume an offline preprocessing step to generate and distribute the precomputed Beaver multiplication triples, OT correlations, and garbled circuits to the two non-colluding clouds. In practice, the preprocessing can be implemented by another independent cloud provider or by the patients participating in the computation. In our benchmarks, we focus on measuring the online cost of the protocol execution in terms of both communication as well as end-to-end protocol execution time. We additionally measure the total size of the precomputed values. Finally, to simulate the two non-colluding cloud servers for the online phase of the computation, we place the two clouds on opposites coasts of the United States, and we measure the total protocol execution time and communication.

## Acknowledgements

We thank members of the Boneh and Bejerano labs for valuable discussions, tools and project feedback. All patient studies performed under Stanford IRB. The DDD study presents independent research commissioned by the Health Innovation Challenge Fund [grant HICF-1009-003], a parallel funding partnership between Wellcome and the Department of Health, and the Wellcome Sanger Institute [grant WT098051]. The views expressed in this publication are those of the authors and not necessarily those of Wellcome or the Department of Health. The study has UK Research Ethics Committee approval (10/H0305/83, granted by the Cambridge South REC, and GEN/284/12 granted by the Republic of Ireland REC). The UK research team acknowledges the support of the National Institute for Health Research, through the Comprehensive Clinical Research Network. This work was supported in part by Stanford Graduate and Computational and Evolutionary Human Genomics Fellowships (K.A.J.), NSF Graduate Research Fellowship (D.J.W.), Stanford Interdisciplinary Graduate Fellowship (J.B.), Simons and National Science Foundation Fellowships (D.B.), Stanford Pediatrics Department, DARPA, Packard Foundation and Microsoft Faculty Fellowships (G.B.). The authors declare no conflict of interest.

## Supplementary Figures

**Supplementary Figure 1.**
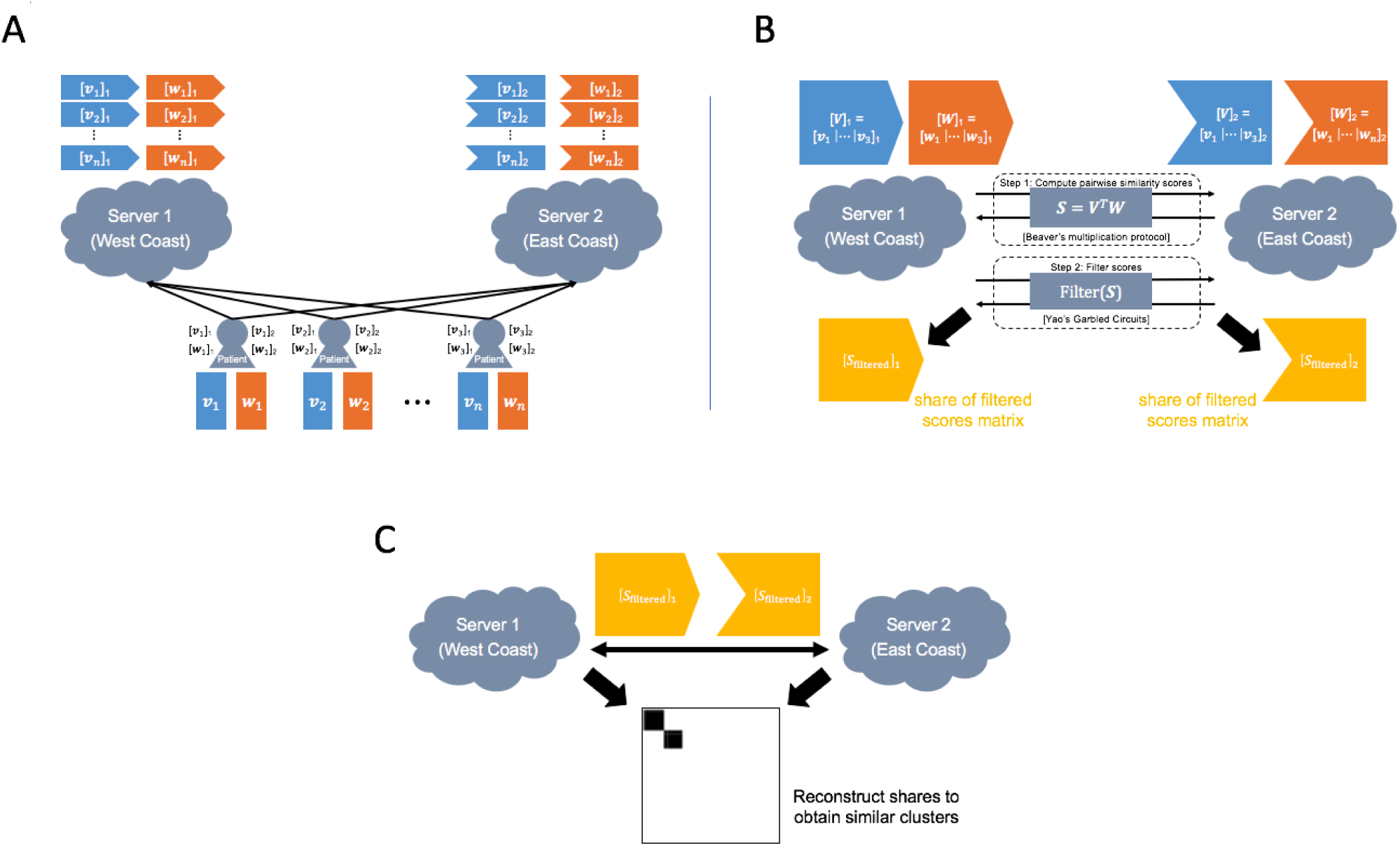
Scenario 1: Patient COHORT BUILDING protocol description. As part of the protocol setup each patient creates two vectors based on their phenotype information: the indicator phenotype vector ***v*** and the weighted phenotype weights ***w***. The shares of each vector are sent to the two clouds and the secure protocol is executed between the two clouds to identify cohorts of phenotypically-similar patients. We use (precomputed) Beaver multiplication triples to compute the Phrank set similarity measure between each pair of patients. Then using Yao’s protocol, we threshold the values in the pairwise similarity matrix, and then filter the resulting values to remove small (spurious) clusters (see Methods for details).

**Supplementary Figure 2.**
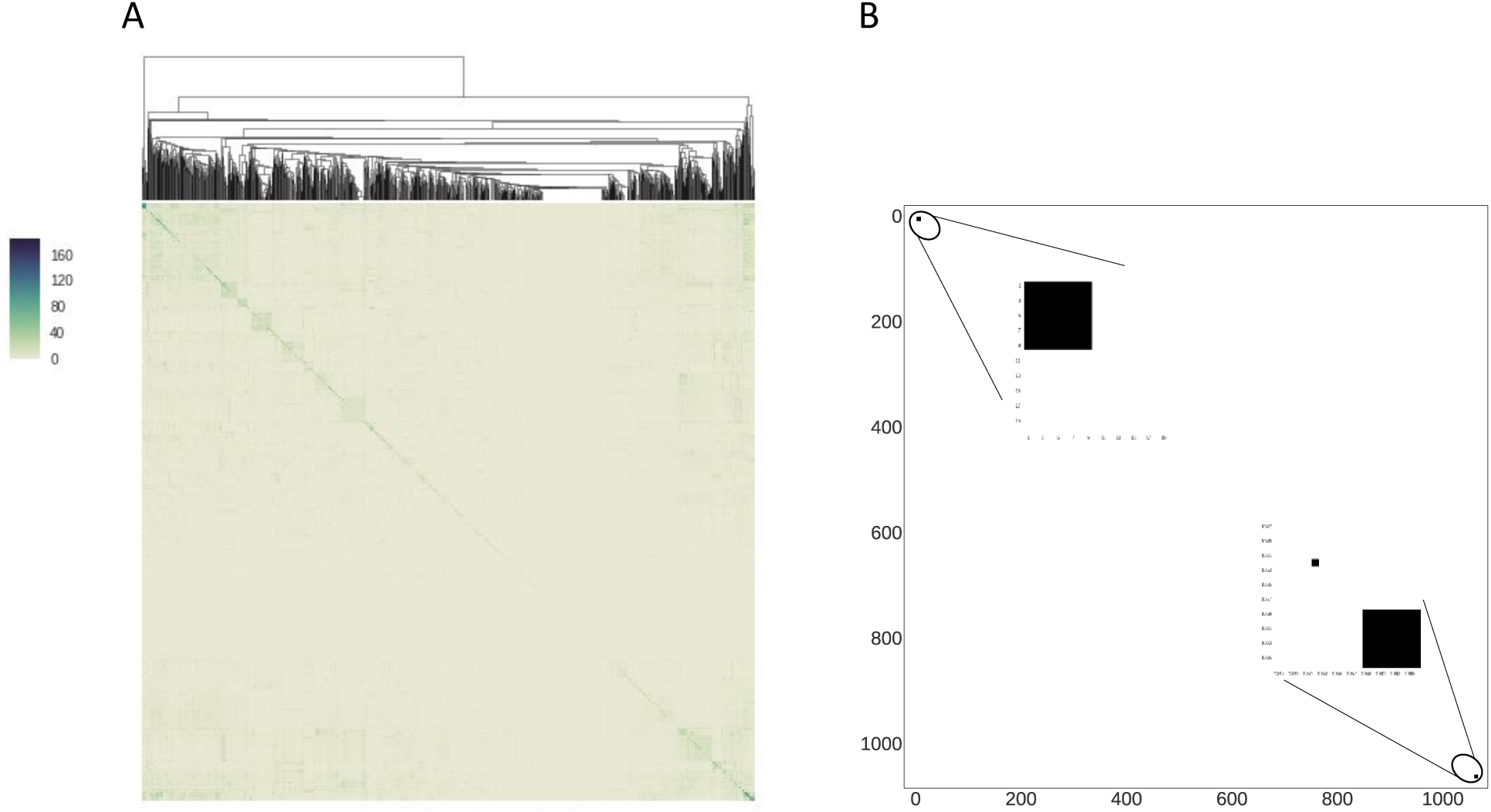
Scenario 1: Small cohort discovery. (A) The pairwise Phrank similarity score matrix for all *n* = 1,069 patients (which no one observes). (B) After clustering and filtering, exactly two cohorts of patients are revealed. Patients are identified by their patient identifiers (while their Phrank scores are completely hidden).

**Supplementary Figure 3.**
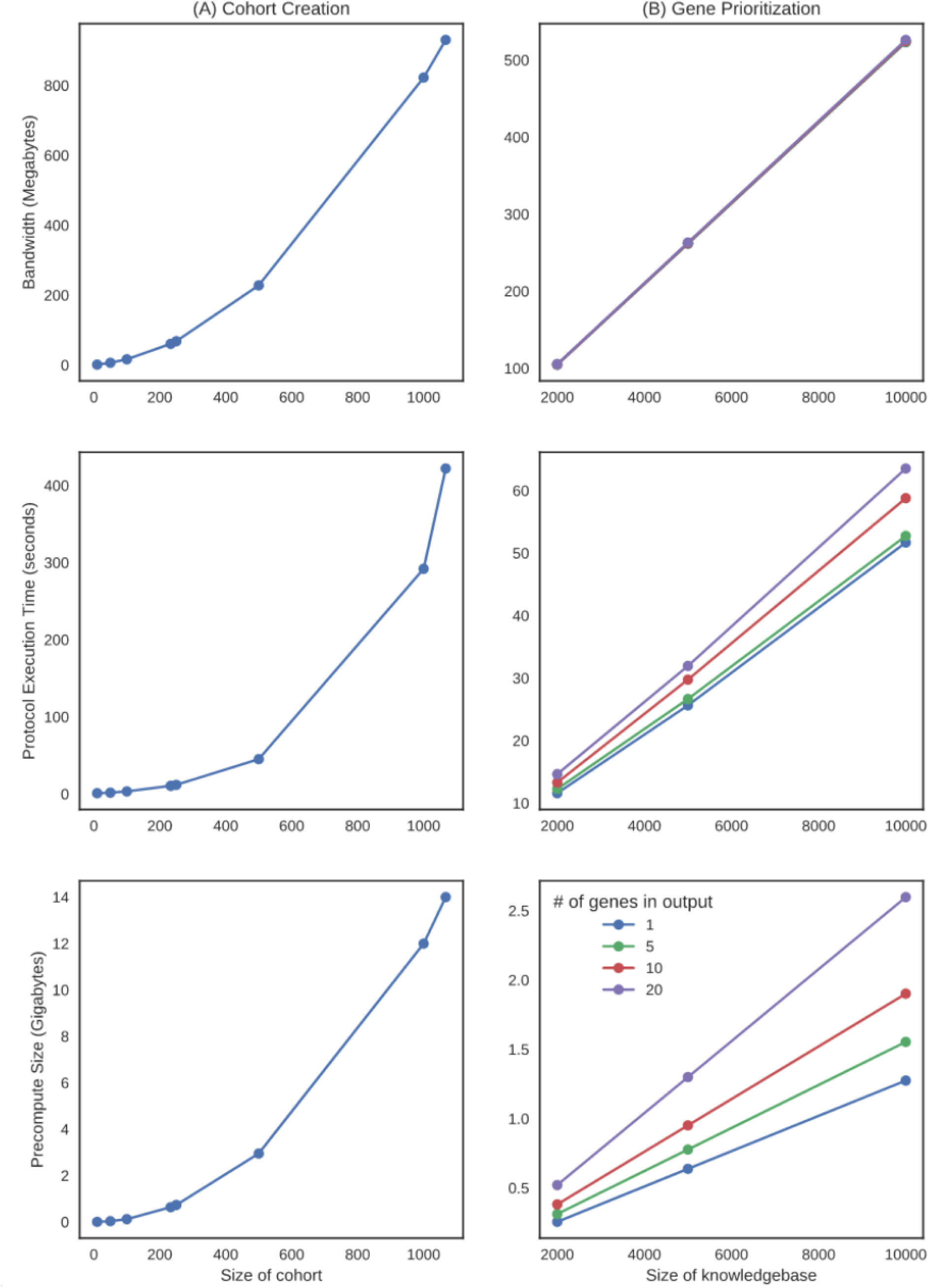
Performance measurements for secure computations. Bandwidth, protocol execution time and the size of the precomputed values for the cohort creation and gene prioritization protocols. All measurements were taken using a single-threaded execution on two Amazon EC2 servers, one located on the East Coast and the other on the West Coast. (A) For the cohort creation protocol, the bandwidth, network time, and precomputation size all scale quadratically with the number of patients. (B) For the gene prioritization protocol, the bandwidth, network time, and precomputation size all scale linearly with the size of the service provider’s knowledgebase. Increasing the number of genes output by the protocol does not have a significant impact on the bandwidth, but does slightly increase the computation time and the amount of required precomputation.

**Supplementary Figure 4.**
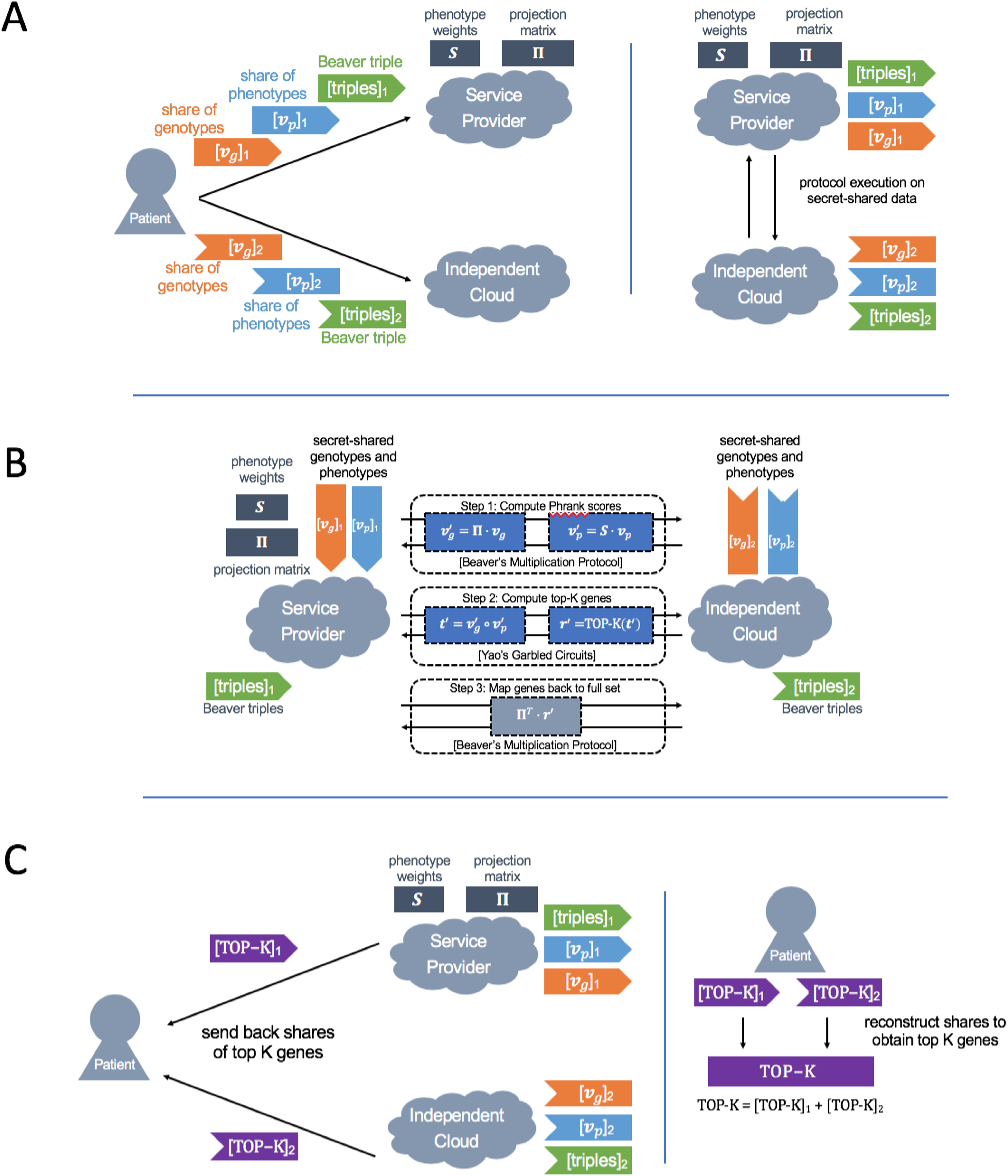
Scenario 2: GENE PRIORITIZATION protocol description. We show how to securely prioritize patient candidate genes based on their likelihood for causing the patient’s disease using patient phenotypic and genotypic information, and without revealing any patient information to the genome analysis provider. The patient begins by securely uploading their genotypic and phenotypic information to a non-colluding cloud, who will facilitate the computation. (A) The gene prioritization protocol is performed between an individual and a third-party genome analysis provider. The individual starts by sharing their data with the non-colluding cloud and the server using a secure secret-sharing protocol. This ensures that neither the service provider nor the non-colluding cloud can reconstruct the patient’s input data. The genome analysis provider and the cloud then execute a secure computation protocol to identify the prioritized list of genes most likely to explain the patient’s disease. (B) In the first step of the computation, the computing parties apply a projection from the set of all candidate disease genes to the set of genes actually present in the genome analysis provider’s knowledgebase (which is often a smaller subset of genes). This significantly reduces the computational cost of subsequent operations in the protocol. Then the two parties compute the Phrank score between each of the patient’s phenotypes and the phenotypes associated with each gene in the provider’s knowledgebase. Both of these computations rely on Beaver’s protocol (with precomputed Beaver multiplication triples). Next, using Yao’s garbled circuits, the two parties jointly identify the top 10 genes with the highest Phrank scores. The result is a secret-shared binary vector indicating the top 10 genes. Finally, the two parties project the indices of identified genes back into the set of all possible genes. This is the output of the protocol. This step again relies on Beaver’s multiplication protocol. (C) At the end of the protocol, the two clouds send back their shares of the output to the client, who adds them together to learn the top 10 genes most correlated with their phenotypes.

